# Benthic composition drives functional composition of fish assemblages despite varying recreational diving intensity

**DOI:** 10.64898/2026.07.28.741252

**Authors:** Sophie Porton, Alba Iana McClinton, Leah Brinch-Iversen, Caitlin Elizabeth Bolton, Alexander McMaster, Irini Valanto Papageorgiou, Megan Mary Soulsby, Miguel Barbosa

## Abstract

Human activities are driving widespread degradation of natural ecosystems, with coral reefs among the most affected. While the impacts of climate change and pollution on coral reefs are well documented, the ecological consequences of recreational scuba diving, a rapidly growing industry, remain less understood. Namely, few studies have explored how diving influences trait diversity of coral reef fish assemblages, a key dimension of ecosystem resilience. Here, we investigated spatial variation in benthic composition and fish trait diversity across reef sites in Utila, Honduras, an island heavily reliant on dive tourism. We surveyed five sites of varying levels of diving intensity to quantify benthic cover and fish assemblages. Contrary to expectations, sites with high diving intensity did not show reduced coral cover, with the most heavily dived site (Stingray Point) maintaining the highest overall coral cover. Algae, however, dominated the benthos across all sites. While functional richness was maintained across the diving intensity gradient, functional composition shifted significantly among sites. This compositional shift was driven primarily by the Von Bertalanffy growth coefficient (k). Sites with low hard coral cover but higher benthic heterogeneity harboured fish communities dominated by fast-growing, early maturing species, whereas high hard coral cover sites supported assemblages of slower-growing, later-maturing species. Benthic composition, rather than diving intensity, emerged as the primary filter of fish life-history strategies. These findings suggest that while current diving intensity may not erode the volume of functional trait space, it is associated with distinct benthic states that filter for specific life-history strategies. Our results highlight the importance of maintaining benthic heterogeneity in reef systems, even in the absence of obvious coral loss. As functional diversity underpins key ecosystem services, it is important that coral reef monitoring focuses on changes in trait composition rather than just simpler taxonomic diversity metrics. Achieving this in collaboration with dive operators to promote low-impact diving is essential to sustaining reef resilience in tourism-dependent regions like Utila.

## INTRODUCTION

Coral reefs provide a disproportionate number of ecosystem services due to their exceptional biodiversity and intricate structural complexity (1, 2). This complexity underpins a wealth of ecosystem services, including vital support for fisheries, effective coastal protection, and substantial tourism revenue, with coral reef services estimated to generate US$352,000 per hectare annually (3). Despite their economic and ecological importance, coral reefs worldwide face unprecedented challenges from diverse human activities (1, 4, 5), with these changes particularly impacting developing nations heavily reliant on reef-derived benefits (6, 7). Understanding the intricate ways in which human impacts affect coral reef functioning and biodiversity is paramount for effective conservation and the preservation of their ecological, social, and economic roles. This study addresses this need by focusing on one such impact: recreational diving on coral reefs.

The vast diversity and clear, warm waters make coral reefs highly attractive for tourism, generating an estimated US$36 billion globally each year (8). For small Caribbean nations, coral reef tourism related activities generate US$7.9 billion in revenue per year, representing around 10% of the GDP in the region (9). However, this economic benefit comes with ecological costs. There is accumulating evidence indicating that recreational divers can negatively affect coral reef health and functionality through both direct and indirect physical damage (10-13). Direct damage can occur via divers or their equipment trampling or touching corals, leading to breakage and reduced habitat availability for reef organisms (14, 15). Indirect damage can arise from boat anchors or from the resuspension of sediment by fin kicks, which can potentially suffocate coral polyps (16, 17). While individual diver impacts may seem minor, their cumulative effects can be detrimental to vulnerable and slow-growing reefs (10, 16). Furthermore, diver presence can alter fish behavior, such as reducing grazing activity and potentially leading to macroalgal overgrowth (18, 19).

Despite the documented physical impacts of recreational diving on corals, the relationship between recreational diving and the functional diversity of associated fish assemblages remains less studied. Functional diversity, which links species’ traits to ecosystem processes (20), offers a stronger approach to understanding the ecological ramifications of biodiversity change (21-24). Here we address this gap by examining the impact of recreational diving on benthic composition and the functional diversity of associated fish assemblage, in Utila, Honduras. By focusing on functional traits, this study therefore provides a novel and potentially more insightful perspective on the ecological effects of recreational diving on coral reef ecosystems. Utila has benefitted greatly from recreational diving tourism since the early 1990s, with this small island now hosting roughly 16 dive centres that regularly cater to divers of all levels (25, 26). In fact, an estimated 85% of the island’s economy is sustained, directly and indirectly by the dive industry (25). However, this high economic reliance on the dive industry makes Utila especially vulnerable to the degradation of the surrounding coral reef functioning, making their conservation and preservation a crucial priority for the island.

Recreational diving can alter benthic composition through physical contact, contributing to shifts in the relative cover of hard coral (13, 27, 28). These changes in benthic state matter ecologically because the composition of the benthos determines the availability and diversity of habitats for associated fish communities (29-31). Reefs dominated by structurally complex hard coral support a different set of ecological niches (i.e. more refugia, more foraging microhabitats, more stable conditions) than reefs where coral has been replaced by algae, rubble, or sand (32). This shift in habitat availability is expected to filter for different fish life-history strategies. For example, hard coral rich benthic states tend to support slow-growing, habitat-specialist species, whereas more homogenised or algae-dominated states favour opportunistic, fast-maturing species better able to exploit variable or disturbed conditions (30, 32). Therefore, rather than tracking species identities alone, the functional diversity metrics used here can reveal whether these ecological roles and processes are maintained or eroded as benthic composition changes (20, 22). The mechanistic link, from diving related structural benthic composition changes to shifts in habitat availability, to filtering of fish life-history strategies provides the underlying framework for testing two main predictions. First, sites with higher recreational diving intensity will show reduced coral cover and greater benthic homogenisation. Second, benthic homogenization will be associated with a shift in fish functional composition toward fast-growing, opportunistic life-history strategies, and a corresponding reduction in functional richness. We also consider the important caveat that dive operators may preferentially select naturally resilient sites, which we discuss in the context of interpreting our results.

## MATERIAL & METHODS

### Study area

This study was conducted along the southern coast of Utila (16°06′N, 86°54′W), one of the Bay Islands of Honduras. Utila is situated approximately 30 km north of the Honduran mainland and forms a critical component of the Mesoamerican Barrier Reef System, the second largest barrier reef in the world (33).

Five distinct dive sites with comparable reef topography and depth profiles were selected for this study. All sites were located on the southern side of Utila (Figure 1) and were categorized based on their primary mode of dive access and expected diving intensity: moored sites, unmoored sites, and a shore dive site. This categorization facilitated the comparison of benthic composition and fish diversity across varying levels of recreational diving pressure.

**Figure 1.**
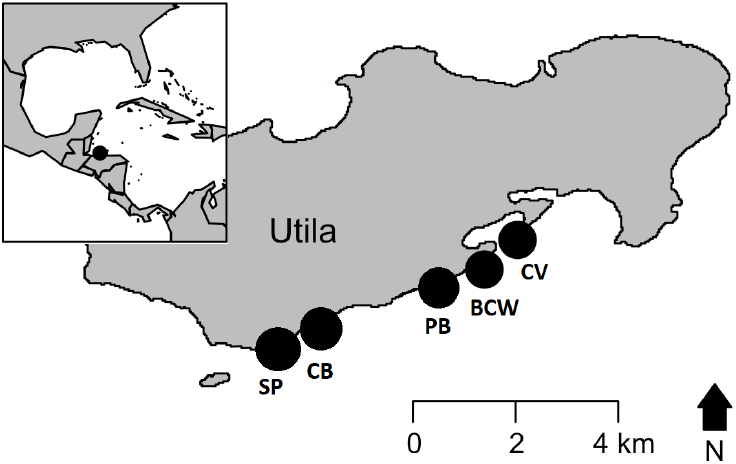
Map showing the location of the island of Utila (Honduras) with the five reef study sites indicated. Reef site names are abbreviated as: Black Coral Wall (BCW), Cabañas (CB), Coral View (CV), Pretty Bush (PB) and Stingray Point (SP).

Moored Sites (High Diving Intensity): Black Coral Wall (BCW) and Stingray Point (SP) are popular dive sites equipped with permanent moorings. These sites experience consistently high levels of recreational diving activity from numerous local dive shops and Operation Wallacea expeditions.

Unmoored Sites (Lower Diving Intensity): Pretty Bush (PB) and Cabanas (CB) were previously equipped with moorings, which were removed approximately two years prior to data collection (March 2018). This removal was assumed to have resulted in a reduction in dive boat traffic and, consequently, a lower intensity of recreational diving compared to moored sites.

Shore Dive Site (Variable Diving Intensity): Coral View Resort (CV) is a shore-accessible dive site directly adjacent to the Coral View Resort. This site experiences high diving and snorkelling activity, particularly during the summer months, primarily from Operation Wallacea participants and resort guests. Unlike the moored sites, Coral View experiences minimal boat traffic or usage from other external dive operators.

Information regarding scuba diving intensity at each site was gathered through informal interviews conducted with multiple dive shop owners on Utila and staff members of Operation Wallacea, an environmental research and conservation expedition organization with a permanent base at Coral View Resort on Utila.

The south coast reef line consists of a spur and groove system (34) and all selected sites were within a few hundred meters of each other, ensuring broadly comparable environmental conditions despite their distinct habitat characteristics. High and low diving intensity sites were selected in pairs based on proximity along the coast, allowing usage intensity to be compared between otherwise similar, nearby sites.

### Benthic and fish surveys

Data were collected during March 2020 using a standardized transect-based survey method, adapted from the Reef Life sampling protocol (35). The survey was employed to simultaneously estimate benthic composition and fish assemblage at each of the five study sites. At each dive site, three 50m transects were laid parallel to the reef slope at a constant depth of 12m. Transects were meticulously deployed to ensure no overlap and maintain spatial independence while allowing for continuous sampling coverage within the designated area. Diver roles were rotated between dives to mitigate observer bias.

### Benthic composition

Benthic composition was quantified using randomized quadrat sampling along each transect line. Ten 1m^2^quadrats were deployed and photographed at randomly generated positions along each 50m transect, with the constraint that at least one quadrat was placed within every 5m interval. This sampling procedure has been shown to provide an adequate representation of benthic cover comparable to other established methods (36). Photographs of the quadrats were subsequently analysed using the image processing software ImageJ (37). For each quadrat image, the area covered by different benthic components was manually delineated and categorized using the free-snip tool. The area of each categorized component was then automatically calculated by the software. The benthic categories used for quantification were: Hard coral, Algae, Soft coral/Sponges, and Other benthos, which included abiotic substratum such as sand, uncolonized reef rock, and coral rubble. Live and dead coral were merged into a single benthic category, as both statuses provide habitat for fish species (38). Finally, we recorded the morphology of each hard coral colony observed during benthic surveys, classifying colonies into eight growth forms: arborescent, columnar, digitate, encrusting, foliose, massive, tabular, and branching (39).

### Fish assemblage

Underwater Visual Surveys (UVS) were conducted to quantify the fish assemblage, using a strip transect method (40). Fish data were collected by a diver swimming slowly along each 50m transect line, identifying and recording every fish encountered within a 2m strip width (S1). An inventory list of frequently observed species was utilized to enhance the efficiency and accuracy of underwater record-keeping. While it is acknowledged that diver presence can influence fish behavior (e.g., attraction or deterrence of certain species), UVS are widely considered an adequate and reliable method for describing fish communities (41). To minimize potential disturbance to fish, the diver assigned to fish census data consistently swam ahead of the diver responsible for deploying benthic quadrats.

## STATISTICAL ANALYSIS

### Benthic composition

To investigate differences in benthic community structure among sites, benthic composition was analysed at two levels: at the overall benthic category level (hard coral, algae, soft coral/sponges, and other benthos) and, within the hard coral level, the coral growth form level (arborescent, columnar, digitate, encrusting, foliose, massive, tabular, and rod). This two-level approach allowed us to determine whether sites differed in the broad composition of the benthos and, if so, whether those differences reflected changes in the structural architecture of the coral community or simply in the total cover of coral relative to other benthic categories.

Differences among sites were tested using permutational multivariate analysis of variance (PERMANOVA) using the adonis2 function in the R package *vegan* (42). A Bray-Curtis dissimilarity matrix was constructed from the proportional data. For the benthic category level, proportional cover was averaged across the ten quadrats per dive to avoid pseudo-replication. For the growth form level, proportions were calculated as the number of colonies of each growth form divided by the total number of hard coral colonies recorded per dive. The assumption of homogeneity of multivariate dispersions was verified at both levels using the betadisper function, with significance assessed via permutation.

Following the multivariate analysis, differences in the cover of individual benthic categories among sites were examined using a Generalized Linear Model (GLM) with a quasibinomial error distribution. The quasibinomial family was selected to accommodate the proportional nature of the response variable and to correct for overdispersion (43). The model was structured as proportion cover as a function of the interaction between Site (Stingray Point, Black Coral Wall, Cabanas, Pretty Bush, and Coral View) and benthic category. This interaction allowed for the assessment of site-specific differences within each benthic category. Pairwise comparisons between sites for each benthic category were conducted using Estimated Marginal Means (EMMs) using the emmeans package in R, applying Tukey’s adjustment for multiple comparisons.

### Taxonomic and Trait diversity

Species richness and evenness were calculated for each site. Species richness was calculated as the total number of species observed per dive, and species evenness was quantified using Pielou evenness index. Both metrics were estimated using the *Vegan* package (42). Due to the nature of the species richness count data, which often exhibits overdispersion, we fitted a Generalized Linear Model (GLM) with a negative binomial error distribution using the glm.nb function from the *MASS* package (44). To test for differences in richness among sites, we compared the full model against a null model (richness ∼ 1) using a Likelihood Ratio Test (LRT) with a Chi-squared distribution. Species evenness was modelled using a GLM with a Gaussian error distribution. Like for richness, the significance of the site effect was evaluated by comparing the full model to a null model via an Analysis of Variance (ANOVA).

To test for differences in fish trait diversity between sites, we first selected nine fish traits (Supplementary information 1) representing key aspects of ecosystem functioning (21, 23). Trait data were primarily obtained from FishBase (45, 46) using the R package rfishbase (47). For missing trait values (<5% of the matrix), we retained gaps rather than applying imputation, as empirical evidence suggests that such methods can introduce bias (48). Traits were scaled and centred to ensure equal weighting.

We constructed a trait matrix for all 83 fish species and computed pairwise functional dissimilarity using Gower’s distance metric, which accommodates mixed categorical and continuous traits (22). The resulting distance matrix was ordinated via Principal Coordinates Analysis (PCoA) to visualize trait distributions in a reduced-dimensional functional space. We retained 6 PCoA axes, which captured approximately 75% of the total functional variation (quality of representation = 0.74). Following this, we calculated two trait diversity indices for each site: Functional Richness (FRic), defined as the volume of functional space occupied by the community, and Functional Evenness (FEve), defined as the regularity of abundance distribution within the trait space. Differences in FRic and FEve across sites were tested using Generalized Linear Models (GLM). Additionally, differences in overall functional composition were tested using PERMANOVA on the Gower dissimilarity matrix.

All statistical analyses were conducted in R (49).

## RESULTS

### Benthic Composition

Benthic community composition differed significantly across sites (PERMANOVA: F=2.21, p=0.03, R^2^=0.42). Pairwise comparisons show that the benthic composition at site SP (high dived site) was the most distinct, differing significantly from site CV (least dived site) (F=4.62, p=0.022, R^2^=0.43, Figure 2a).

**Figure2a.**
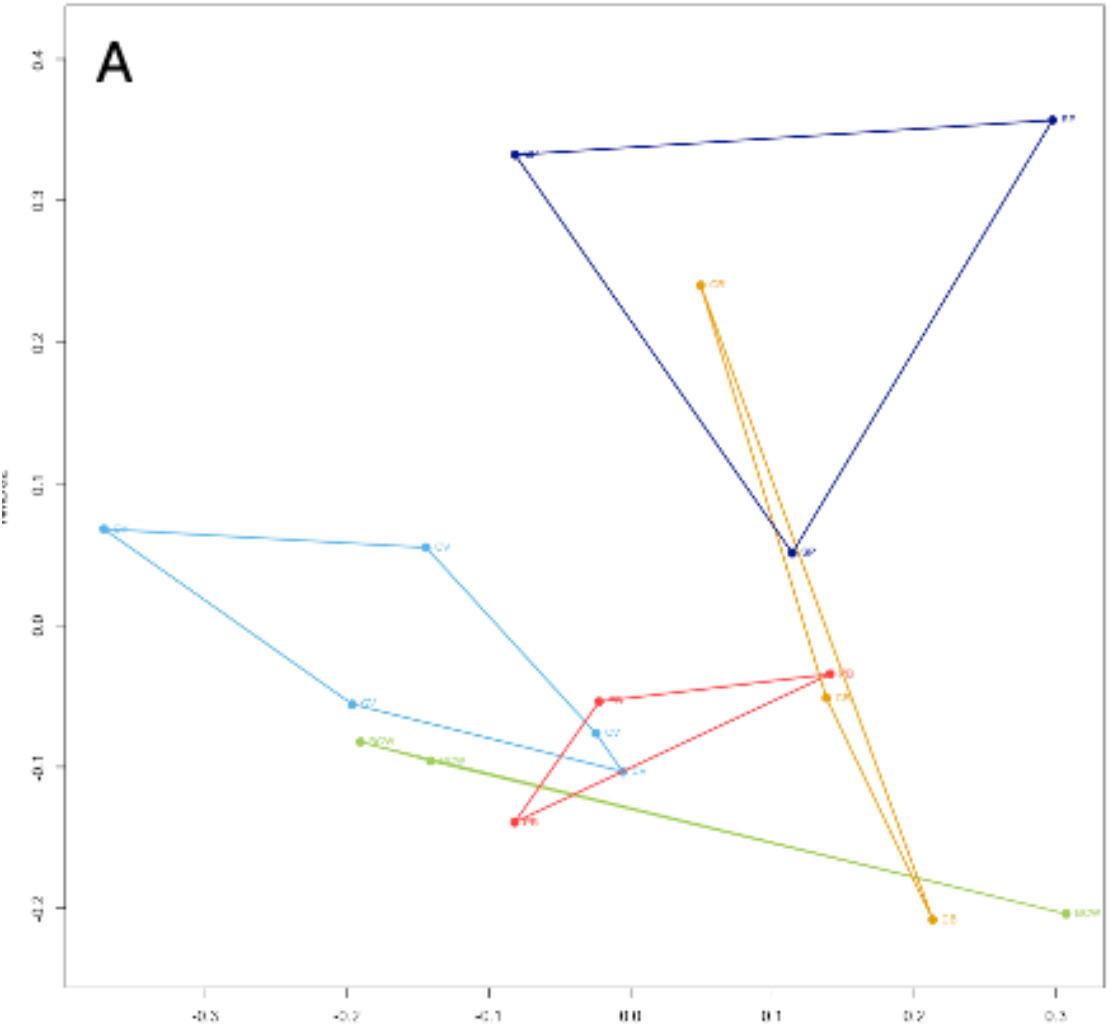
Non-metric multidimensional scaling (nMDS) ordination of Bray-Curtis dissimilarities in benthic cover, showing the spread of dive-level observations (points) for each of the five sites. Each polygon represents the convex hull of replicates within a site. Sites showing greater separation reflect more distinct benthic community composition.

The Generalized Linear Model (GLM) revealed a significant interaction between site and benthic composition (GLM, F=5.27, p<0.001). Algae was the dominant benthic category across all sites, exhibiting the highest proportion of cover (Figure 2b, Table 1). While Algae dominated, SP (high diving site) exhibited the highest cover of hard coral, which was significantly greater than that observed at BCW (p = 0.036), PB (p=0.009) and CV (p<0.001). Algae cover at BCW was significantly higher than at SP (p=0.022). There were no significant differences in the cover of soft coral/sponge among the sites.

**Figure2b.**
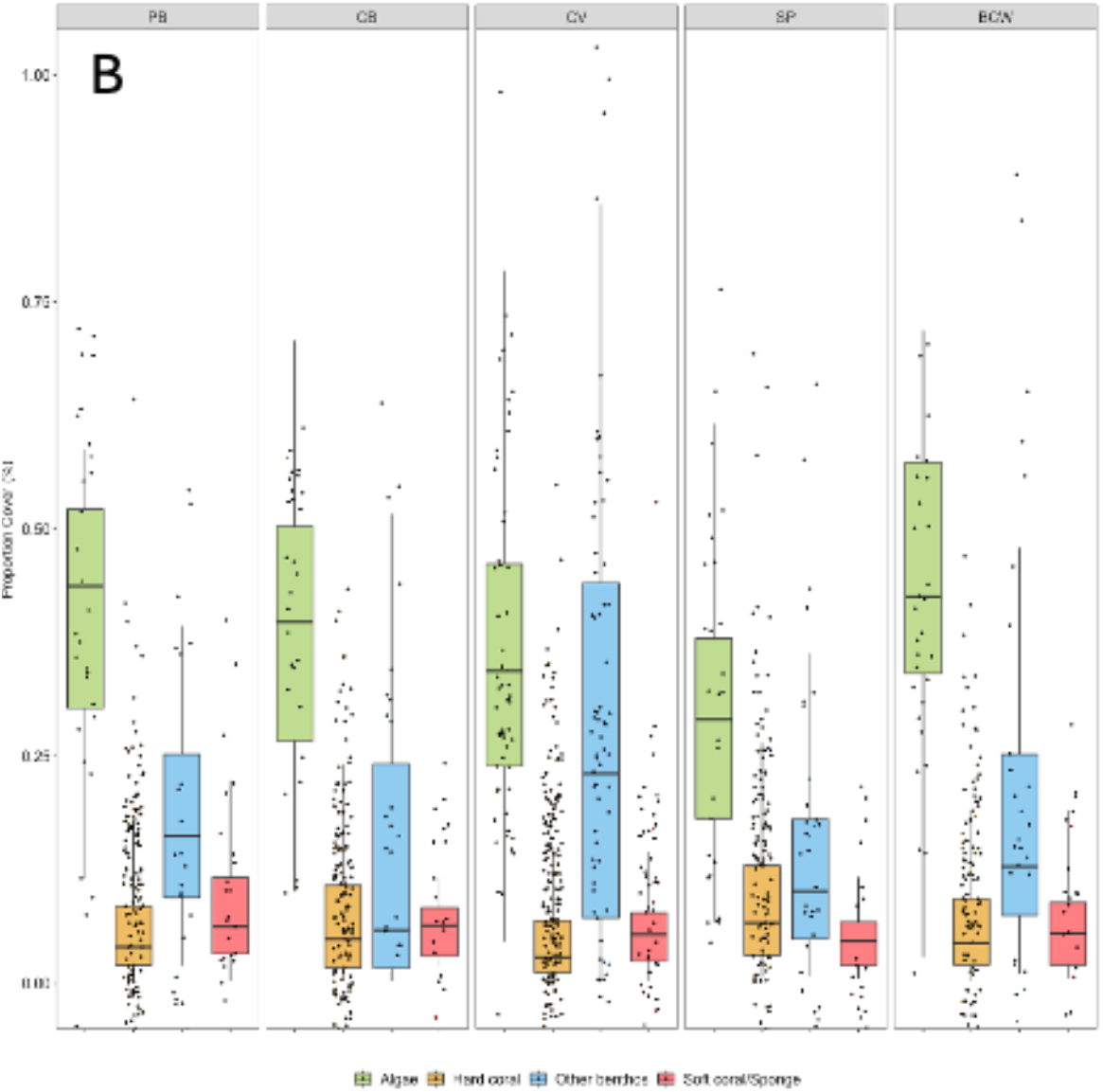
Proportional cover (%) of four benthic categories — algae, hard coral, other benthos and soft coral/sponge, across the five reef sites. Error bars denote 1.5× the IQR. Raw data shown

**TABLE 1.** proportion of benthic categories across study sites.

| Sites | Algae | Coral | Other benthos | Soft coral/Sponge |
| --- | --- | --- | --- | --- |
| BCW | 0.4317706 | 0.07029020 | 0.2332365 | 0.06567113 |
| CB | 0.3984234 | 0.07422696 | 0.1409048 | 0.06428338 |
| CV | 0.3546740 | 0.06286223 | 0.2841825 | 0.06455184 |
| PB | 0.4051583 | 0.06818372 | 0.1906947 | 0.08276644 |
| SP | 0.2975854 | 0.10473327 | 0.1496176 | 0.04884045 |

Coral growth form composition did not differ significantly among sites (PERMANOVA: F = 0.82, p = 0.58, R^2^ = 0.035). All sites were broadly similar in the relative representation of the eight growth forms recorded, with massive and encrusting corals the most consistently abundant morphologies across sites (Figure 2c).

**Figure2c.**
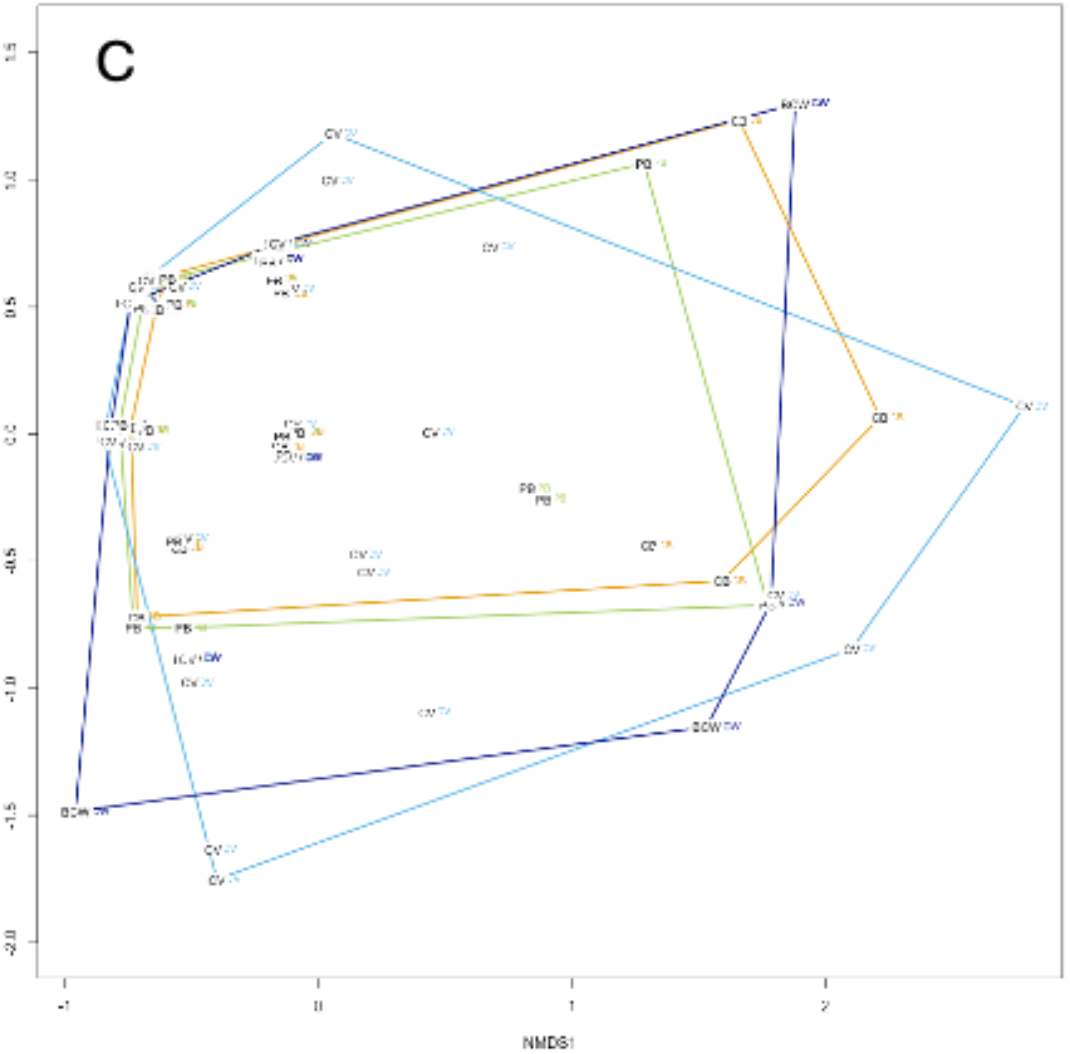
Non-metric multidimensional scaling (nMDS) ordination of Bray-Curtis dissimilarities in coral growth forms, showing the spread of dive-level observations (points) for each of the five sites. Each polygon represents the convex hull of replicates within a site. Clear overlap across all sites which denotes no clear distinction in the composition of growth forms across sites

### Taxonomic Diversity

There were no significant differences in species richness across sites (Likelihood Ratio Test, χ^2^ = 3.211, p=0.523). The two heavily dived sites (BCW and SP) exhibited a trend towards lower mean species richness compared to the other sites (BCW – 22.2 species, SP – 23.9 species), with pairwise comparisons showing no significant differences (BCW vs SP, p=0.890) (Figure 3a). Similarly, the low diving pressure sites (PB and CB) and the variable pressure diving site (CV) showed no significant differences in mean species richness (Figure 3a). The estimated mean species richness was lowest at BCW (22.2 species) and SP (23.9 species), and higher at PB (31.7 species), CV (28.8 species), and CB (28.9 species, Figure 3a).

**Figure 3.**
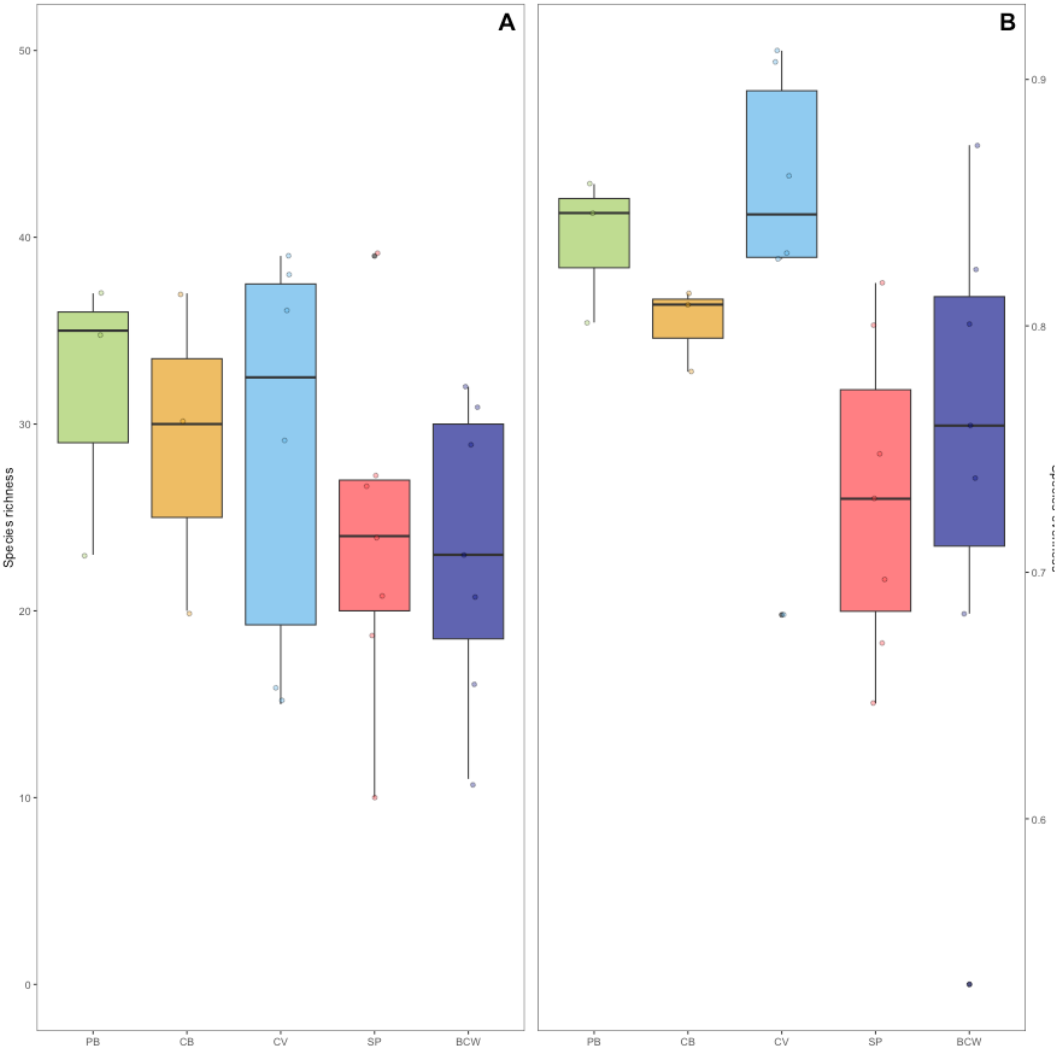
Taxonomic diversity of reef fish assemblages across the five reef sites. (A) Species richness (total number of species recorded per dive transect) and (B) Species evenness (Pielou’s J index) per dive transects for each site. Error bars denote 1.5× the IQR. Raw data shown.

There were no significant differences in species evenness across sites (Likelihood Ratio Test, χ^2^ = 0.055, p=0.072). The variable dived site (CV) and the least dived site (PB) showed a trend towards higher mean evenness compared to the other sites (CV =0.84, p = 0.053; PB=0.83, p = 0.119) (Figure 3b). Conversely, the heavily dived sites (BCW and SP) showed slightly lower, but statistically similar, mean evenness values (0.74 and 0.73, respectively, Figure 3b).

### Trait Diversity

Fish trait composition differed significantly across sites (PERMANOVA: F=1.84, p=0.04, R^2^=0.26, Figure 4). However, there were no differences in functional richness (p=0.21) and functional evenness (p=0.11) between sites (Figure 5). Analysis of Community Weighted Means revealed that this shift was largely driven by differences in the von Bertalanffy growth coefficient (k) between sites (p<0.001, Figure 6). The shore access site with variable diving intensity (CV) had the highest mean value of k (0.64), whereas SP (high intensity diving) exhibited the lowest k value (0.48).

**Figure 4.**
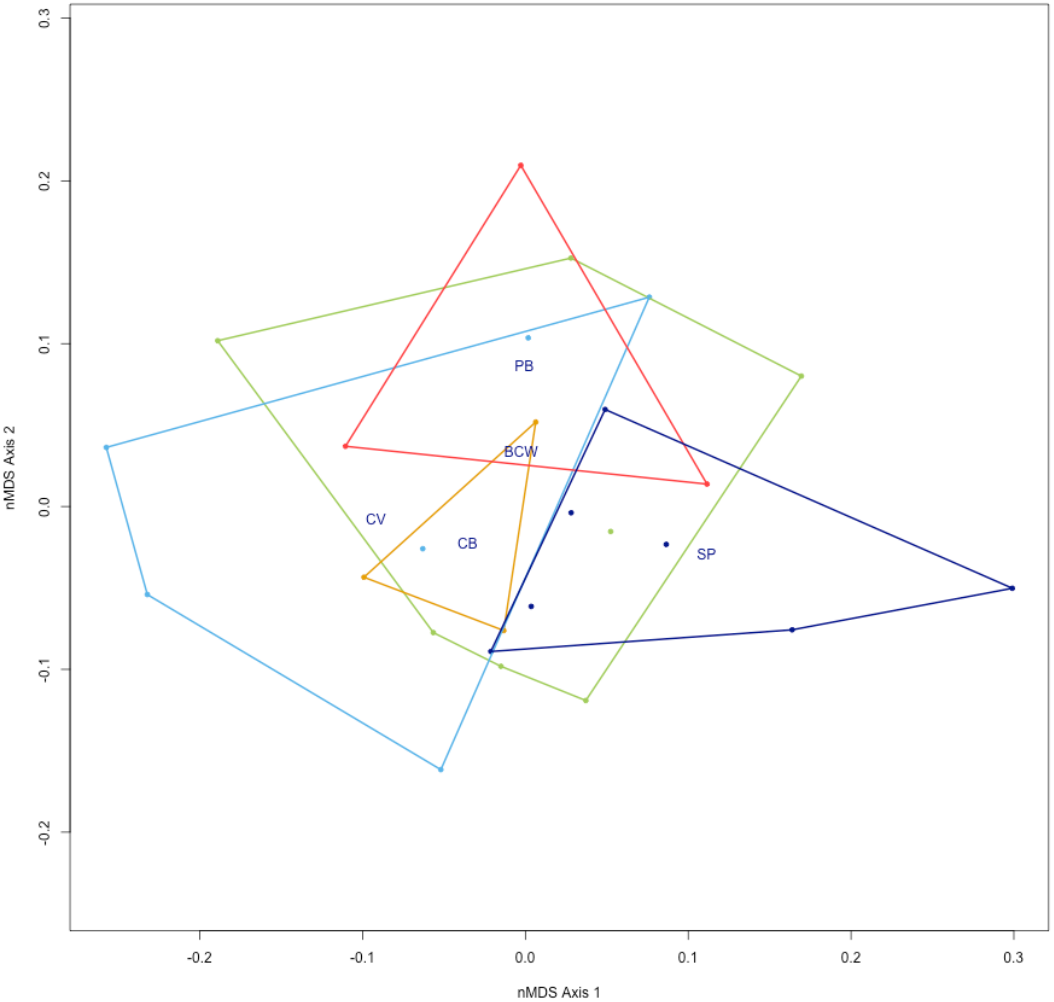
Non-metric multidimensional scaling (nMDS) ordination of Gower dissimilarity in functional trait composition, showing individual dive-level observations and convex hulls for each site. Greater separation between site hulls reflects greater differences in the identity of functional traits represented

**Figure 5.**
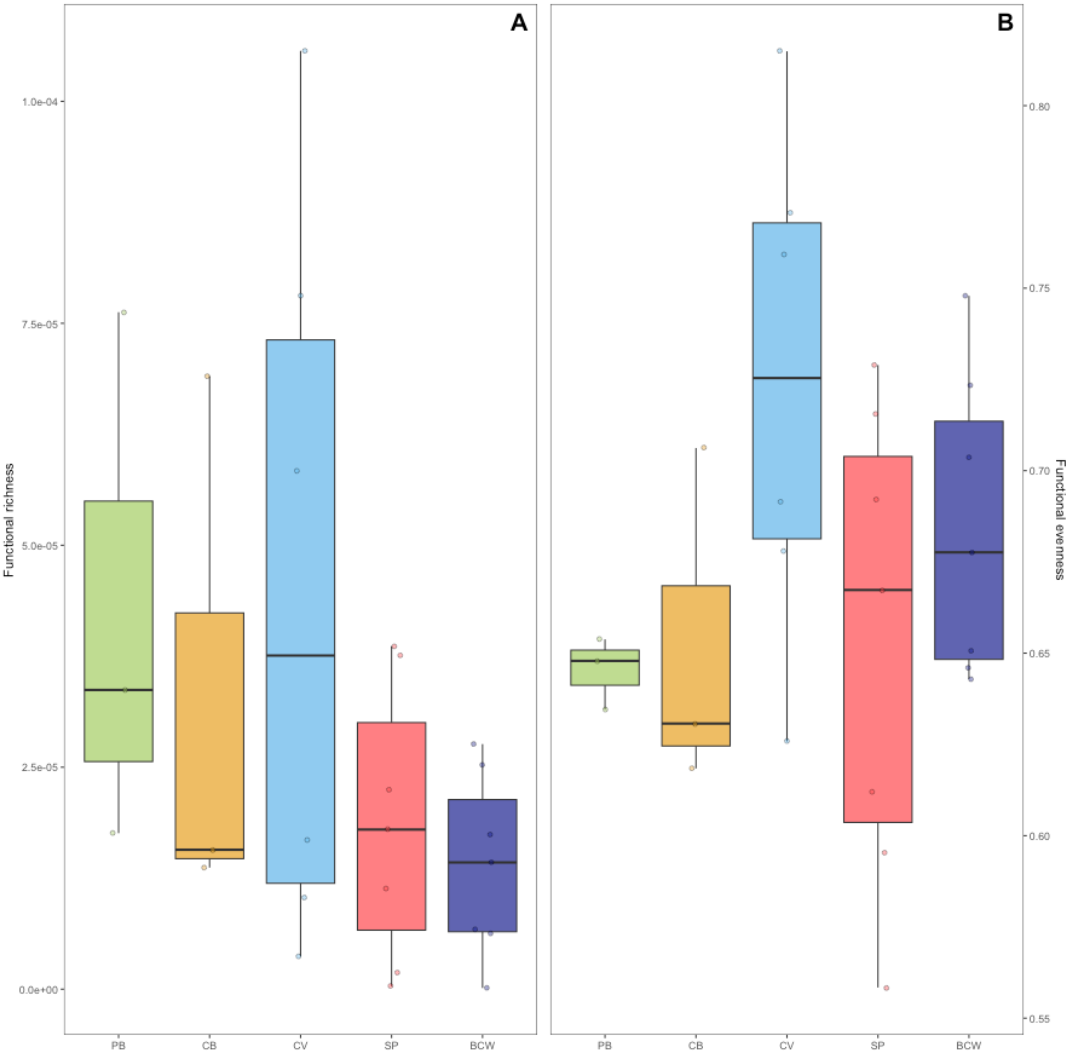
Functional diversity of reef fish assemblages across the five reef sites. (A) Functional Richness and (B) Functional Evenness of fish assemblages at each site. Functional Richness represents the volume of functional trait space occupied by the assemblage, whereas Functional Evenness represents the regularity of species distribution within that space. Error bars denote 1.5× the interquartile range. Range of raw values shown.

**Figure 6.**
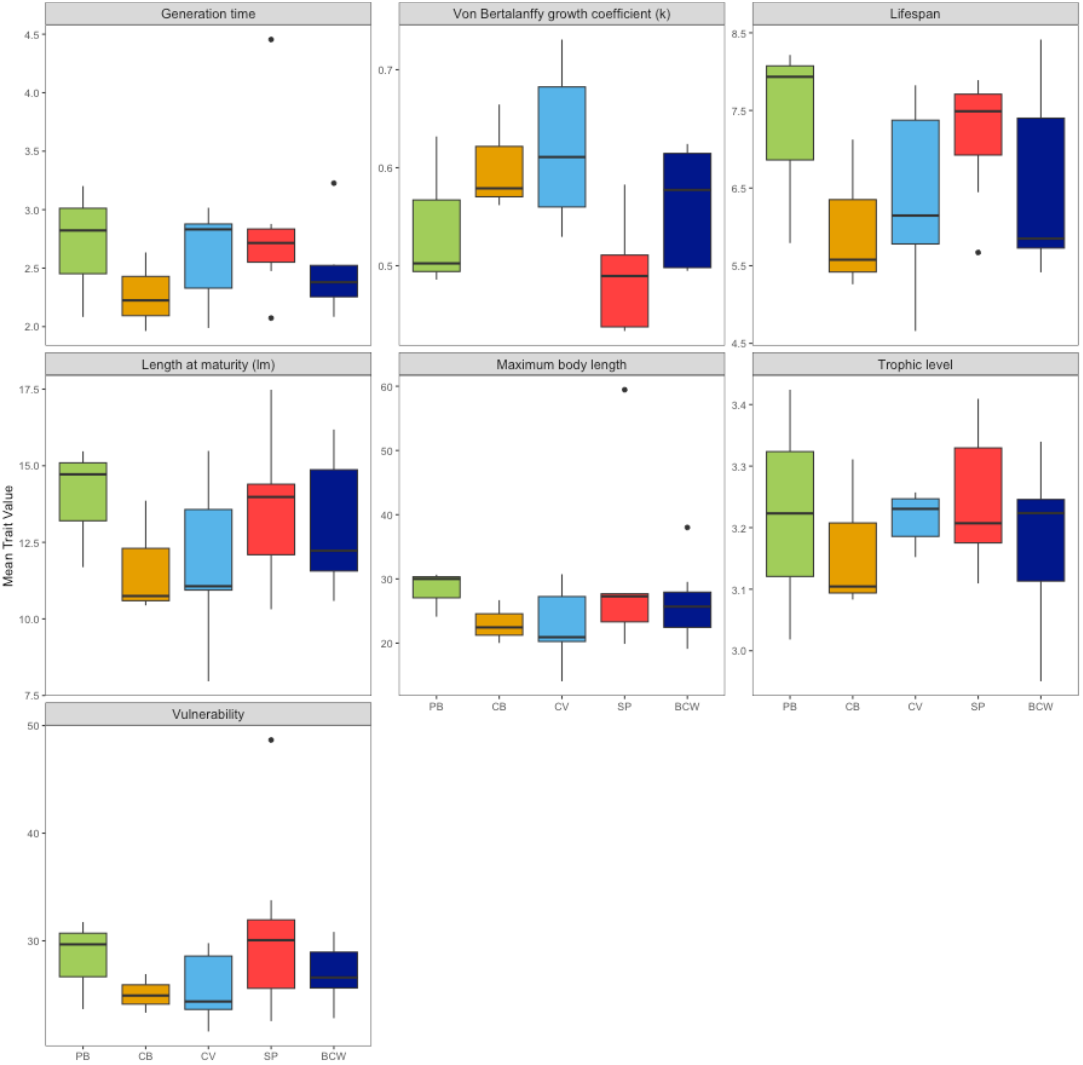
Community Weighted Mean (CWM) values for the functional traits used to characterized functional diversity in fish assemblage across the study sites, per dive transect. Traits include, Generation time, von Bertalanffy growth coefficient (k), Lifespan, Length at maturity (lm), Maximum body length, Trophic level, and Vulnerability

## DISCUSSION

Functional diversity in fish assemblages was broadly maintained across sites, with no significant differences detected in Functional Richness (FRic) or Functional Evenness (FEve). This trait stability suggests a degree of functional redundancy within the reef system (50), where the volume of functional trait space occupied remains similar despite site-level environmental variation. This pattern is consistent with broader evidence from marine systems that species richness is a poor predictor of ecosystem functioning (51). Large-scale observations of fish communities across northwestern European seas, found that ecosystem functioning was unrelated to species richness but was significantly associated with community evenness, species composition, and abiotic variables, a finding that mirrors our results at the reef scale.

While statistical differences in FRic were not detected, Coral View the site with variable diving pressure and the most balanced benthic composition, exhibited the highest functional richness, whereas Black Coral Wall (high diving pressure) showed the lowest. These trends align with the significant differences observed in functional composition, indicating that while the volume of functional space is constant, the identity of traits defining that space shifts between sites. The continuity and diversity in benthic categories at Coral View site provides a wider array of niches and microhabitats for fish (29, 32). This result partially supports the prediction that less homogenized reefs support more functionally diverse fish assemblages but challenges the hypothesis that diving intensity is the primary driver of this diversity.

The absence of a clear link between diving intensity and functional richness raises an important question about causality: are diving companies selecting sites based on inherent ecological diversity, or are their activities shaping it? It is possible that sites like Stingray Point were selected for heavy use precisely because of their inherent resilience and high coral cover, allowing them to sustain high diving pressure without functional degradation. Conversely, sites with homogenous benthos might be avoided by operators or visited less frequently, confounding the "diving intensity” gradient with pre-existing habitat quality. This "selection bias” is a critical consideration in tourism impact studies. It suggests that the resilience of functional richness at high-use sites may reflect the robustness of the initial benthios rather than the absence of diving impacts. Future studies should aim to compare sites with similar initial ecological potential but differing management regimes to disentangle these effects. From a management standpoint, this selection bias also implies a practical opportunity: if operators already gravitate toward structurally robust sites, formalising this tendency through site zoning or rotation policies could reduce cumulative pressure on the most ecologically vulnerable reefs and help maintain the benthic heterogeneity that, as our results show, underpins functional diversity.

A key driver of this compositional shift was the von Bertalanffy growth coefficient (k), which differed significantly across sites. Coral View (site of variable diving pressure) was characterized by a higher mean value indicating a community dominated by species with fast life histories (i.e., rapid growth, early maturation, and shorter lifespans). In contrast, Stingray Point (site of high diving intensity) exhibited a lower mean value (0.48), reflecting a dominance of slower-growing, later-maturing species. This divergence likely reflects underlying habitat stability. Stingray Point maintained the highest hard coral cover (47.5%), a habitat typically associated with long-lived, specialist species (52). Conversely, Coral View had the lowest hard coral cover but the highest cover of "Other Benthos” (e.g., sand, rubble). The prevalence of fast-growing fish species at Coral View can be adaptive to more heterogeneous sites (32). High hard coral cover sites offer greater spatial heterogeneity, more refugia, and richer trophic resources, which sustain slow-growing specialist species that require stable, complex habitat to complete their life cycles (29, 52). Conversely, sites dominated by sand, rubble, and algae offer fewer of these resources, favouring generalist species capable of rapid growth and early reproduction that can exploit transient or disturbed conditions (53). The k gradient we observed therefore most plausibly reflects this benthic filtering process rather than a direct behavioural response to diver presence. This filtering of life-history strategies mirrors patterns observed in other ecosystems under anthropogenic pressure. For instance, in temperate freshwater systems, stream reaches subjected to hydrological variability or physical disturbance often exhibit a shift toward "fast” life-history strategists (r-selected species), whereas stable, pristine headwaters retain "slow” strategists (K-selected species) (54). Similarly, in terrestrial forestry, logged forests often see a rapid turnover toward fast-growing generalist birds and mammals, while old-growth forests retain slow-reproducing specialists (55). In our study, the dominance of slow-growing species at the heavily dived Stingray Point suggests that high diving intensity per se does not automatically induce a shift toward "fast” life histories. Instead, the benthic habitat (high coral cover) appears to be the stronger filter, sustaining slow-growing species despite frequent human visitation. This finding nuances our understanding of diving impacts, by suggesting that the physical degradation of the reef structure (loss of coral), rather than the mere presence of divers, is the critical trigger for shifts in fish life-history composition.

Interestingly, hard coral cover did not correlate with functional richness. Coral View had the lowest live coral, yet the richest functional assemblage. On the other hand, Stingray Point had the highest coral cover but ranked second in functional richness. This result supports the view that reef functionality is preserved even with reduced coral cover, provided structural heterogeneity remains (30, 56). On the other side of the spectrum, we have Black Coral Wall (heavily dived site) which exhibited a trend of both the lowest functional richness and the highest algal cover. This site lacked large carnivorous and herbivore species present at other sites. The loss of functional diversity can disproportionately reduce ecosystem function and resilience (57-59). This is consistent with evidence from large-scale marine fish surveys, which found that reduced ecosystem functioning was associated with assemblages dominated by generalist species, with the loss of high trophic level consumers (51). This result aligns with the compositional shift we observe at Black Coral Wall. The loss of large herbivores also explains the high percentage of algae at Black Coral Wall and the potential state shift (60). As a heavily dived site, Black Coral Wall may experience elevated physiological stress on corals (61). This physiological stress may, over time, lead to reduced coral cover, consistent with evidence that reducing recreational diving pressure can increase coral cover (62). Notably, coral growth form composition did not differ significantly among sites, indicating that sites were broadly similar in the relative representation of structural morphologies. This suggests that the observed differences in functional composition are driven by variation in the overall cover of coral relative to other benthic categories, rather than by selective loss of particular coral morphologies. Such result, emphasises that benthic state as a whole, rather than coral architecture alone, is the primary filter of fish life-history strategies. This highlights the need for precautionary approaches to managing anthropogenic impacts.

Contrary to our hypothesis, there was no consistent relationship between recreational diving intensity and either benthic composition or fish trait diversity. High-use dive sites spanned the spectrum of ecological states: Stingray Point was relatively hard coral rich with slow-growing fish assemblages, while Black Coral Wall was algae-rich with lower functional richness. Coral View, despite at times experiencing high recreational diving intensity, supported a functionally rich assemblage, possibly explained by its shore-based access, which reduces boat-related stressors. The lack of a clear recreational diving impact suggests that local management practices, such as mooring use or site rotation, may be mitigating direct damage, or that diving impacts are mediated by specific coral morphologies rather than just overall coral cover (17, 63). For instance, Stingray Point’s relatively high hard coral cover persisted despite high recreational diving use, perhaps due to the dominance of robust morphologies. Future studies should examine diving intensity in relation to coral morphology. We suspect that branching corals would be more susceptible to diver impact than massive or plate-like forms (63). Alternatively, this pattern may partly reflect selection bias, whereby operators preferentially use sites that were already structurally resilient. Additionally, while diver presence can alter fish behavior (19), the variation in functional richness among sites in our study, rather than a consistent reduction, suggests no systematic deterrence effect. Sites with low recreational diving activity (e.g., Cabanas and Pretty Bush) had intermediate functional richness, while high-use sites spanned both ends of the richness spectrum. Boat traffic, often correlated with diver pressure, may further influence benthic condition and fish trait composition and should be measured independently.

## CONCLUSION

Recreational diving is a major economic player for Utila. Our findings suggest that, at current levels, it may not be causing significant functional degradation of the reef ecosystem. However, we also show that functional composition, specifically the life-history strategies of fish assemblages, is more sensitive to the state of the benthos than to diving intensity alone. Where structural complexity is reduced, whether through recreational diving, bleaching, or other stressors, the functional identity of reef fish communities shifts in predictable ways. This has a direct implication for monitoring and management. As our results highlights, tracking changes in trait composition, rather than species counts alone, provides an earlier and more ecologically meaningful signal of reef degradation (56). For sites dependent of recreational diving, this means combining best diving practises with trait-based reef monitoring to detect functional shifts. It is important to understand that reefs can appear taxonomically intact while undergoing meaningful functional change, and only by looking beyond species richness can managers and operators ensure the long-term resilience of coral reef systems and their services.

